# Cdc42 promotes Bgs1 recruitment for septum synthesis and glucanase localization for cell separation during cytokinesis in fission yeast

**DOI:** 10.1101/754390

**Authors:** Udo N. Onwubiko, Julie Robinson, Rose Albu Mustaf, Maitreyi E. Das

**Affiliations:** Department of Biochemistry & Cellular and Molecular Biology, University of Tennessee, Knoxville, TN, USA

**Keywords:** GTPase, Cdc42, cytokinesis, Bgs1, Eng1, Agn1

## Abstract

Cytokinesis in fission yeast involves actomyosin ring constriction concurrent to septum synthesis followed by septum digestion resulting in cell separation. A recent report indicates that endocytosis is required for septum synthesis and cell separation. The conserved GTPase Cdc42 is required for membrane trafficking and promotes endocytosis. Cdc42 is activated by Guanine nucleotide exchange factors (GEFs). Cdc42 GEFs have been shown to promote timely initiation of septum synthesis and proper septum morphology. Here we show that Cdc42 promotes the recruitment of the major primary septum synthesizing enzyme Bgs1 and consequent ring constriction. Cdc42 is also required for proper localization of the septum digesting glucanases at the division site. Thus, Cdc42 is required to promote multiple steps during cytokinesis.

## Introduction

Cytokinesis involves the separation of a mother cell into two new daughters. In most eukaryotic cells with the exception of plant cells, cytokinesis initiates with the assembly of a medially positioned actomyosin ring that constricts in a regulated manner to form the cleavage furrow [1, 2, 3]. In the fission yeast *Schizosaccharomyces pombe*, in addition to a medially positioned contractile actomyosin ring, cytokinesis requires the synthesis of a tri-layered cell wall structure called the division septum [4]. Once assembled, the actomyosin undergoes a maturation period where a plethora of proteins necessary for cytokinesis are recruited [5, 6, 7]. Upon completion of maturation, septum synthesis initiates concomitantly with ring constriction. The division septum is a complex structure, consisting of the primary septum which is primarily synthesized by the linear-β-glucan synthase Bgs1, and secondary septum, which is synthesized mainly by the branched B-glucan synthase Bgs4 [8, 9, 10]. Once ring constriction completes, the septum matures, and its middle layer (primary septum) gets degraded by digestive enzymes Eng1 and Agn1 to separate the two new daughter cells, thus completing cytokinesis [11, 12]. Improper regulation of cytokinesis could result in non-viable cells via cell lysis, or stalled cytokinesis.

The small GTPase is a master regulator of cell polarity in fission yeast and has been shown to regulate membrane trafficking [13, 14, 15, 16]. We have previously shown that the small GTPase Cdc42 is activated at the division site in a sequential manner to promote timely initiation of septum ingression and septum formation [17]. Cdc42 is activated by the Guanidine nucleotide exchange factors (GEFs) Gef1 and Scd1 [18, 19]. During cytokinesis, loss of Gef1 delays the initiation of ring constriction and Bgs1 recruitment, and loss of Scd1 results in aberrant septum morphology [17]. It is unclear however whether these roles are specific to the GEFs or Cdc42 activity. Cdc42 has also been implicated in the specific trafficking of Bgs1 containing vesicles to the sites of polarized growth, although the mechanism is not known [16]. Gef1 has also been shown to promote endocytosis and proper organization of the scaffold protein Cdc15 [20].

In this study, we assess the specific role(s) of Cdc42 during cytokinesis using a temperature sensitive Cdc42 allele -*cdc42-1625* [15, 21]. We find that disruption of Cdc42 activity resulted in defective cytokinesis, impaired Bgs1 localization to the division site, and disrupted localization of the septum remodeling enzymes Eng1 and Agn1. Our results define specific roles of Cdc42 during cytokinesis. We demonstrate that Cdc42 is required for ring constriction by ensuring Bgs1 recruitment and cell separation by ensuring proper localization of Eng1 and Agn1.

## Results and Discussion

### *cdc42-1625* mutants display similar septation index with *cdc42*+ cells

To investigate the direct role of Cdc42 in cytokinesis we used the temperature-sensitive Cdc42 allele *cdc42-1625*, to disrupt Cdc42 activity at the non-permissive temperature of 36°C. Both *cdc42*^+^ and *cdc42-1625* cells were first grown at the permissive temperature of 25°C, then shifted to 36°C for 4 hours as previously performed. We observed that *cdc42-1625* mutants appeared wider than *cdc42*^+^ cells even at 25°C, which became further pronounced and often caused cell death at 36°C (Fig. 1A). This observation agrees with previous reports that the cdc42*-1625* allele is hypomorphic even at the permissive temperature of 25°C [21]. We analyzed the number of septated cells in each of the experiments, and find that while *cdc42*^+^cells displayed an average septation index of ∼17% at 25°C, and ∼15% at 36°C, *cdc42-1625* strains had septation indices of ∼22% at 25°C and ∼23% at 36°C (Fig. 1B). Although we see a mild increase in the septation index both at permissive and non-permissive temperature, the difference is not statistically significant (Fig. 1B).

**Figure 1.**
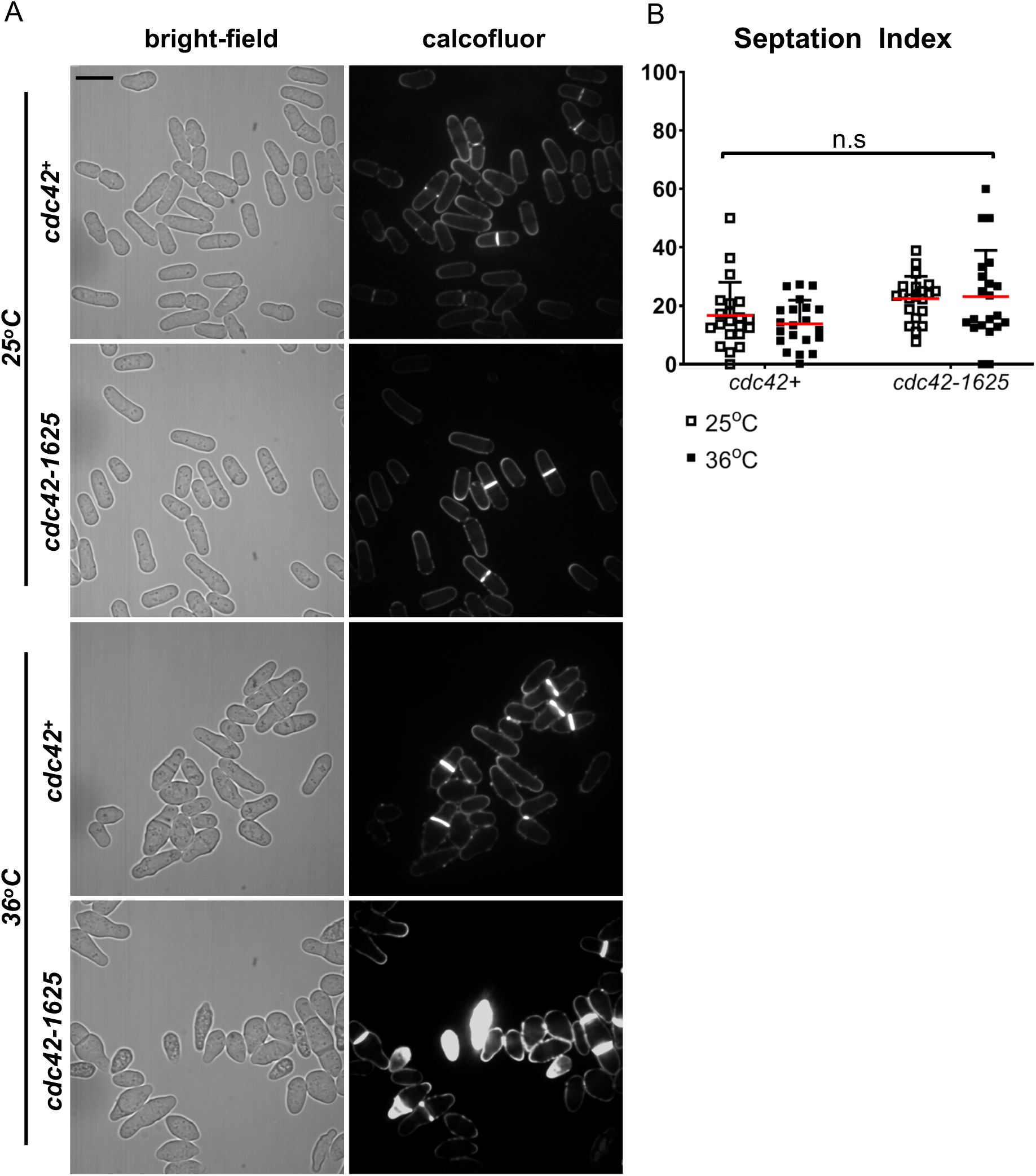
*cdc42-1625* mutants display similar septation index with *cdc42*+ cells. **A.** Representative images of *cdc42*^*+*^ and *cdc42-1625* at permissive (25°C) and non-permissive (36°C) temperatures; [Scale bar: 10µm]. **B.** Septation indices for *cdc42*^*+*^ and *cdc42-1625* at permissive (25°C) and non-permissive (36°C) temperatures. Each point represents the septation index for one image. [Red lines on graph represent the mean; n.s, not statistically significant].

### Cdc42 is required for Bgs1 recruitment to the medial ring during cytokinesis

To investigate why the septation index of *cdc42-ts* mutants at 25°C and 36°C was similar to that of *cdc42*^+^ cells (Fig. 1A), we asked whether these mutant cells displayed fewer septated cells due to an inability recruit the major primary septum synthesizing enzyme Bgs1 and initiate septum synthesis. Bgs1 is recruited to the division site after ring assembly, during maturation [17, 22]. We repeated our experiments in *cdc42*^+^ and *cdc42-1625* cells expressing Rlc1-tdTomato as a ring marker, and GFP-Bgs1. We compared GFP-Bgs1 levels in non-constricting *cdc42*^+^ and *cdc42-1625* cells at 25°C and 36°C. We quantified the fluorescence intensities and find that GFP-Bgs1 levels are significantly reduced in *cdc42-1625* mutants at 25°C and 36°C (p<0.0001), compared to *cdc42*^+^ cells (Fig. 2B). Thus, we find that irrespective of the shift in temperature, the temperature sensitive mutation is effective enough to disrupt Bgs1 recruitment. We did not observe any differences in GFP-Bgs1 levels in *cdc42*^+^ cells at 25°C or 36°C (Fig. 2B). We further assessed the number of cells with actomyosin rings containing Bgs1 in *cdc42*^+^ and *cdc42-1625* cells. We find that ∼85% of *cdc42*^+^ cells display actomyosin rings with GFP-Bgs1 at 25°C and 36°C (Fig. 2C). Interestingly ∼89% of *cdc42-1625* cells at 25°C, displayed GFP-Bgs1 containing actomyosin rings, and this value was comparable to *cdc42*^+^ cells at 25°C and 36°C (Fig. 2C). At 36°C however, the percentage of GFP-Bgs1 containing actomyosin rings was significantly reduced to ∼64%. We find that at 36°C, fewer actomyosin rings in *cdc42-1625* cells recruit Bgs1. Cdc42 is therefore required for recruitment of Bgs1 to the division site after ring assembly, which further enables the timely initiation of ring constriction.

**Figure 2.**
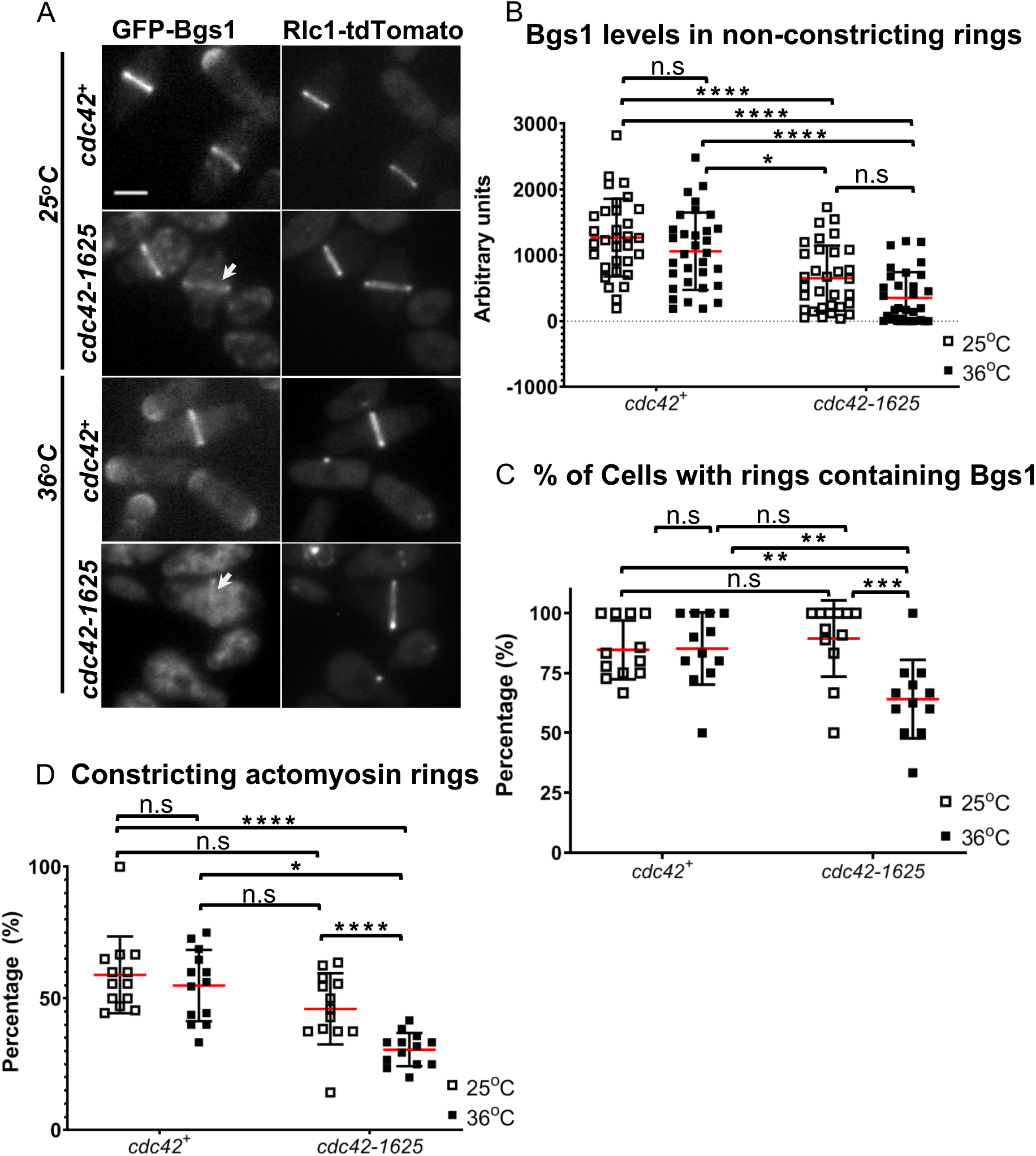
Cdc42 is required for Bgs1 recruitment to the medial ring during cytokinesis. **A.** Sum z-projections of images displaying Bgs1-GFP and Rlc1-tdTomato at permissive (25°C) and non-permissive (36°C) temperatures in control cells and *cdc42-1625* mutants. White arrows point to Bgs1-GFP localization at the ring in cells with non-constricting rings. **B.** Quantification of Bgs1-GFP levels in cells with non-constricting rings in the mentioned strains at permissive and non-permissive temperatures, n=32 cells**. C.** Quantification of the percentage of cells with ring showing Bgs1 localization, n>100 cells, per strain. **D.** Quantification of the percentage of constricting rings observed among cells displaying an actomyosin ring in the mentioned strains, at permissive and non-permissive temperatures. [Red lines on graph represent mean, blue line represents median, for all graphs, *P≤0.05; **P≤0.01; ****P≤0.0001; n.s., not statistically significant; Scale bar: 5µm].

Next, we probed the percentage of *cdc42-1625* mutants that are able to initiate successful ring constriction. We analyzed the percentage of cells showing constricting rings and found no significant differences in *cdc42*^+^ cells at 25°C and 36°C (Fig. 2D). In *cdc42*^+^ cells at 25°C ∼59% of the actomyosin rings appear to be under constriction, while this value is 55% at 36°C. In *cdc42-1625* mutants, the percentage of constricting rings at 25°C, was 47%, but decreased to 31%, after 4 hours at 36°C (Fig. 2D). Our data therefore highlight the importance of Cdc42 for Bgs1 recruitment and initiation of ring constriction in cytokinesis.

### Cdc42 is required for proper localization of digestive enzymes during cell separation

While our data indicated that Cdc42 is required for Bgs1 recruitment to the division site, resulting in fewer constricting rings, it fails to explain why the *cdc42-1625* mutants have a septation index similar to the control cells. A recent report indicated that Gef1 has a role in cell separation [23]. Thus, it is possible that the septation index observed in *cdc42-1625* cells is due to a digestion defect of the septa that formed under permissive conditions. To test this, we assessed the role of Cdc42 in septum digestion that leads to cell separation. Cdc42 has also been implicated in endocytosis [20] and polarized trafficking of the exocyst, which is also spatially positioned at the division site during ring constriction and cell separation [24, 25, 26]. To probe the role of Cdc42 in cell separation, we assessed the localization of the septum digesting enzymes Eng1 and Agn1 in *cdc42*^+^ and *cdc42-1625* strains. Eng1-GFP and Agn1-GFP were simultaneously expressed in *cdc42*^+^ and *cdc42-1625* strains. Cells were grown at the permissive temperature of 25°C, and imaged to assess Eng1-GFP and Agn1-GFP localization. We observed that these enzymes only localized to septated cells (Fig. 3A), and there were no obvious differences in the fluorescence intensities in both *cdc42*^+^ and *cdc42-1625* cells. Next, we generated 3D-reconstructions of Eng1-GFP Agn1-GFP localization at the division site, to assess localization patterns. We first looked in *cdc42*^+^ cells and found three different localization patterns: a ring, a ring and dot, and a disc-like localization (Fig. 2B (i-iii)). For proper septum digestion these enzymes need to be delivered to the outer rim of the membrane barrier in a ring like pattern [25].

**Fig. 3.**
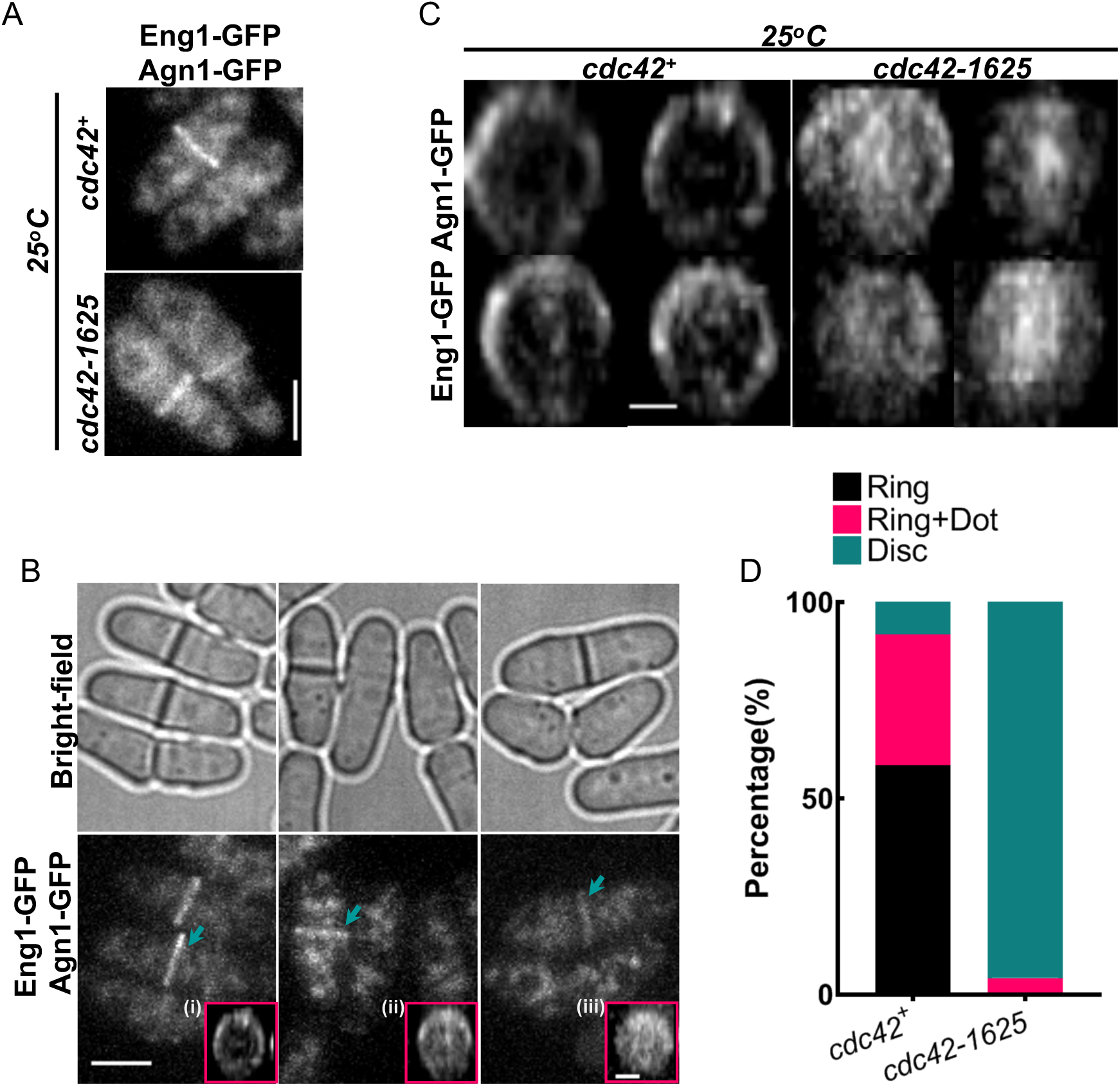
Cdc42 is required for proper localization of digestive enzymes during cell separation. **A.** Sum z-projections of images displaying GFP tagged digestive enzymes Eng1 and Agn1 at permissive temperature (25°C) in control cells and *cdc42-1625* mutants, scale bar: 5µm. **B.** Bright-field and fluorescent images of Eng1-GFP and Agn1-GFP in control cells undergoing cell separation. The pink boxes highlight the 3d-reconstructions of the different localization patterns of Eng1-GFP and Agn1-GFP observed: (i) Ring, (ii) Ring + Dot, (iii) Disk. **C.** 3D-reconstructed division site of Eng1-GFP Agn1-GFP localization at the division site during cell separation in the strains listed, at 25°C, Scale bar: 5µm. **D.** Quantification of Eng1-GFP and Agn1-GFP localization patterns at the division site in the strains mentioned. Scale bar for all 3D-reconstructed division site: 2µm.

3D-reconstructions of Eng1-GFP Agn1-GFP in *cdc42*^+^ cells mainly displayed ‘ring’ and ‘ring and dot’ localization patterns. In *cdc42-1625* mutants at 25 °C, Eng1-GFP Agn1-GFP mostly displayed the ‘disc-like’ pattern (Fig. 3C), suggesting that these enzymes were not able to localize properly. We quantified the frequency of these patterns within each strain and find that 58% of *cdc42*^+^ cells, display rings, 33% display a ‘ring +dot’ and 8% display ‘disc-like’ localization. In *cdc42-1625* mutants, 0% display any rings, 4% display a ‘ring +dot’ and 96% display ‘disc-like’ localization (Fig. 3D). We were unable to observe the enzymes in cells grown at 35°C. The localization of the digestive enzymes Eng1 and Agn1 is thus not polarized, rather appears very diffuse at the division site in *cdc42-1625* cells, suggesting that Cdc42 is required to properly localize these enzymes to enable proper septum digestion.

## Conclusions

Cytokinesis is a tightly regulated process, and in fission yeast, septum synthesis and ring constriction are properly coordinated to generate viable daughter cells. We report that Cdc42 is required for proper cytokinesis in fission yeast. We demonstrate that functional Cdc42 is required for the recruitment of the primary septum enzyme Bgs1 and for the proper localization of the septum synthesizing enzymes Eng1 and Agn1. Cdc42 is an essential protein and hence here we used a conditional mutant to investigate its role in cytokinesis. It is possible that Cdc42 has an essential role in cytokinesis and this has not been elucidated to date due to the nature of *cdc42* mutants available. Cdc42 has been reported to play major roles in membrane trafficking events [15, 16, 20]. However, Cdc42 specifically promotes recruitment of Bgs1 but is not required for the recruitment of a secondary septum synthesizing enzyme Bgs4 [17]. Thus, it is likely that the role of Cdc42 in cytokinesis is to regulate the recruitment of specific cargo required for distinct steps in cytokinesis. Delivery of Bgs1 and the glucanases occur at spatially regions of the division site. Future research will reveal how Cdc42 promotes precise spatial recruitment of these different enzymes.

## Materials and methods

### Strains and Cell culture

Strains used in this study are listed in Table 1. All *S. pombe* strains used in this study are isogenic to PN972. Unless otherwise mentioned cells were cultured in yeast extract (YE) medium and grown exponentially at 25°C. Genetic manipulations of strains were carried out using standard techniques (Moreno et al., 1991). Cells were grown exponentially for at least 3 rounds of eight generations for each assay. On the day of the experiment, cells were grown to O.D. of 0.1 and a subset of control and *cdc42-1625* cell culture was transferred to another flask, and placed in a 36°C shaking incubator for 4 hours before imaging.

**Table 1.** Strains list.

| Strain | Genotype | Origin |
| --- | --- | --- |
| 972 | *h-* | *P. Nurse* |
| 527 | *h- ade6-704 leu1-32 ura4-D18 his3-D1* | *P. Nurse* |
| 528 | *h+ ade6-704 leu1-32 ura4-D18 his3-D1* | *P. Nurse* |
| PPG5660 | *h+ cdc42-1625-kanMX leu1-32 ura4-D18* | [21] |
| YMD1452 | *bgs1-GFP:leu1+ rlc1-tdTomato- NAT<sup>r</sup> leu1-32 ura4-D18* | *This study* |
| YMD825 | *h+ eng1-GFP agn1-GFP* | *This study* |
| YMD1412 | *cdc42-1625 GFP-bgs1:leu1 rlc1-tdTomato- NAT<sup>r</sup> leu1-32 ade6-M210 ura4-D18* | *This study* |

### Microscopy

Image acquisition was performed at room temperature (23-25°C) with the VT-Hawk 2D-array scanning confocal microscope (Visitech intl., Sunderland, UK) using an Olympus IX-83 inverted microscope with a 100x/numerical aperture 1.49 UAPO lens (Olympus, Tokyo, Japan). Cells were spun down in a mini desk microcentrifuge, mounted directly onto glass slides with a #1.5 coverslip (Fischer Scientific, Waltham, MA) and imaged promptly. Z-series images were acquired with depth interval of 0.4um. All Images were acquired with MetaMorph (Molecular Devices, Sunnyvale, CA). All images were analyzed with Image J (National Institutes of Health, Bethesda, MD).

Statistical analysis was performed using 2-way ANOVA, followed by Tukey’s HSD Post Hoc Test where appropriate. Comparisons between experimental groups were considered significant when p≤0.05.

### Calcofluor Staining

Cells were grown as described earlier in the methods section, and stained in yeast extract media (YE) with 50µg/ml Calcofluor White (M2R Sigma-Aldrich, St. Louis, MO) at room temperature. Both controls and mutant strains were washed with fresh YE liquid 1x, and imaged immediately.

### Image Analysis

Sum projections of acquired z-series images were performed in Image J. We quantified GFP-Bgs1 levels from sum-projections. To do this, we measured the intensity at the division site in cells with non-constricting actomyosin rings, using a box with an area of 330pixel^2^. Background correction was performed with the cytoplasm for each cell.

## Acknowledgements

We thank Pilar Perez and Fred Chang for strains. This work is supported by the National Science foundation (1616495). U.N.O. and J.R.R. were supported by NIH IMSD (R25GM086761) and are currently supported by NSF GRFP (1452154).

